# SARS-CoV-2 accessory proteins ORF3a and ORF7a modulate autophagic flux and Ca^2+^ homeostasis in yeast

**DOI:** 10.1101/2022.12.29.522217

**Authors:** José Luis Garrido-Huarte, Josep Fita-Torró, Rosa Viana, Amparo Pascual-Ahuir, Markus Proft

## Abstract

Virus infection involves the manipulation of key host cell functions by specialized virulence proteins. The SARS-CoV-2 small accessory proteins ORF3a and ORF7a have been implicated in favoring virus replication and spreading by inhibiting the autophagic flux within the host cell. Here, we apply yeast models to gain insights into the physiological functions of both SARS-CoV-2 small ORFs. ORF3a and ORF7a can be stably overexpressed in yeast cells, producing a decrease in cellular fitness. Both proteins show a distinguishable intracellular localization. ORF3a specifically localizes to the vacuolar membrane, whereas ORF7a targets the endoplasmic reticulum. Overexpression of ORF3a and ORF7a leads to the accumulation of Atg8 specific autophagosomes. However, the underlying mechanism is different for each viral protein as assessed by the quantification of the autophagic degradation of Atg8-GFP fusion proteins, which is inhibited by ORF3a and stimulated by ORF7a. Overexpression of both SARS-CoV-2 ORFs decreases cellular fitness upon starvation conditions, where autophagic processes become essential. These data are in agreement with a model where both small ORFs have synergistic functions in stimulating intracellular autophagosome accumulation, ORF3a by inhibiting autophagosome processing at the vacuole and ORF7a by promoting autophagosome formation at the ER. ORF3a has an additional function in Ca^2+^ homeostasis. The overexpression of ORF3a confers calcineurin-dependent Ca^2+^ tolerance and activates a Ca^2+^ sensitive FKS2-luciferase reporter, suggesting a possible ORF3a-mediated Ca^2+^ efflux from the vacuole. Taken together, we show that viral accessory proteins can be functionally investigated in yeast cells and that SARS-CoV-2 ORF3a and ORF7a proteins interfere with autophagosome formation and processing as well as with Ca^2+^ homeostasis from distinct cellular targets.

## Introduction

The severe acute respiratory syndrome coronavirus-2 (SARS-CoV-2) is causing worldwide the ongoing epidemic of coronavirus disease 2019 (COVID-19), posing a strong threat to public health and economic performance globally (Wu et al., 2020; Huang et al., 2020). Despite the expanding knowledge, which has accumulated in the past years about SARS-CoV-2 virology and epidemiology, we still lack an antiviral drug for efficient COVID-19 treatment (Sanders et al., 2020). With the pandemic still unfolding, continued investigation into the characteristically high infectivity and pathogenicity of SARS-CoV-2 is required.

SARS-CoV-2 is a member of the β-coronavirus genus of the Coronaviridae family and contains a large, continuous genome encoded in almost 30 kb of single-stranded RNA including the information for 14 potential open reading frames (ORFs) (Gordon et al., 2020; Zhang and Holmes, 2020; Zmasek et al., 2022). The major part of the 5’
s-terminal region of the genome contains two overlapping ORFs, orf1ab and orf1a, encoding 16 non-structural proteins (NSPs). The smaller 3’-terminus of the SARS-CoV-2 genome contains the information for the spike (S), membrane (M), envelope (E) and nucleocapsid (N) structural proteins and a relatively high number of small accessory proteins (ORF3a, ORF3b, ORF6, ORF7a, ORF7b, ORF8, ORF9b, and ORF10), which are specific for the viral genus. The non-structural proteins form the replicase necessary for genome replication and mRNA synthesis of the virus, while the structural proteins form the virus particle. Importantly, the accessory proteins have distinct functions in modulating the host cell response to maximize the infection process and pathogenesis (Liu et al., 2014; McBride and Fielding, 2012).

Macroautophagy is one important function, which is generally targeted during viral infection in the host cell (Abdoli et al., 2018; Mao et al., 2019; Jordan and Randall, 2012), and specifically by some SARS-CoV-2 accessory proteins (Zhao et al., 2021). Macroautophagy is an evolutionarily conserved homeostatic mechanism of eukaryotic cells, which selectively removes superfluous or damaged organelles and proteins, as well as invasive microbes (Levine and Kroemer, 2008). This process is essential for maintaining cellular homeostasis and starts with the formation of enclosed double membrane vesicles known as autophagosomes in the cytosol. Autophagosomes fuse with lysosomes in metazoan organisms or vacuoles in fungi in order to digest the transported cargo material by lysosomal/vacuolar enzymes. The autophagic flux from autophagosome formation to degradation is a highly regulated and dynamic process, which can be modulated by both the vesicle production and fusion (Nakatogawa, 2020; Lei et al., 2022). In general, the autophagic vesicle transport can have either pro-or anti-viral functions. As an intrinsic defense mechanism of host cells, it can inhibit viral replication by lysosomal degradation of virus particles (Kobayashi et al., 2014; Richetta and Faure, 2013). However, some viruses have developed strategies to hijack the host autophagic system in order to favor their own replication and spreading (Pradel et al., 2020). In fact, it has been shown that β-coronaviruses use the lysosomal trafficking for egression from the host cell (Ghosh et al., 2020), which implies that normal autophagosome-lysosome function has to be interrupted by the virus for efficient spreading (Shroff and Nazarko, 2021; Miller et al., 2020; Koepke et al., 2021). Indeed, SARS-CoV-2 infection causes an accumulation of autophagosomes by specific accessory proteins such as ORF3a or ORF7a (Hayn et al., 2021; Miao et al., 2021). The molecular function of SARS-CoV-2 ORF3a interrupting the autophagic flux has been recently elucidated. ORF3a blocks the fusion of autophagosomes with lysosomes through the evolutionarily conserved homotypic fusion and protein sorting (HOPS) complex (Miao et al., 2021; Zhang et al., 2021; Qu et al., 2021), which seems to favor the lysosomal egression of the virus from the host cell (Chen et al., 2021). Additionally, the determination of the ORF3a structure and its reconstitution in liposomes revealed a potential Ca^2+^ permeable non-selective cation channel activity (Kern et al., 2021), whose relevance in vivo is still to be confirmed. The autophagy related function of SARS-CoV-2 ORF7a is much less known, although a very recent study showed that this accessory protein is able to both stimulate autophagosome formation and to inhibit autophagosome fusion with lysosomes (Hou et al., 2022).

Given that SARS-CoV-2 ORF3a and ORF7a exert at least part of their function via macroautophagy, a homeostatic process conserved from fungi to humans, we aimed in this work to functionally study the overexpression of both accessory viral proteins in the yeast (*Saccharomyces cerevisiae*) model. We find that both proteins are stably overexpressed in yeast causing a moderate growth inhibition. ORF3a specifically localizes to the vacuolar membrane and causes accumulation of Atg8 specific autophagosomes, while inhibiting Atg8 vacuolar degradation. ORF7a specifically localizes to the endoplasmic reticulum, where it seems to stimulate autophagosome formation. ORF3a additionally interferes with intracellular calcium homeostasis, conferring calcineurin dependent Ca^2+^ tolerance and overactivation of calcineurin signalling. Our data suggest that both accessory proteins cooperate in the simultaneous stimulation of autophagosome formation and interruption of vacuolar fusion with the possible participation of vacuolar Ca^2+^ release.

## Materials and Methods

### Yeast strains and growth conditions

*Saccharomyces cerevisiae* strains used in this study are shown in Table 1. Yeast cultures were grown at 28°C in Synthetic Dextrose (SD) or Galactose (SGal) media containing 0.67% yeast nitrogen base with ammonium sulfate and without amino acids, 50mM succinic acid (pH 5.5) and 2% of the respective sugar. According to the auxotrophies of each strain, methionine (10 mg/l), histidine (10 mg/l), leucine (10 mg/l) or uracil (25 mg/l) were supplemented. Starvation conditions were induced by SD-N medium containing 0.17% yeast nitrogen base without amino acids or ammonium sulfate and 2% glucose. Yeast cells were transformed by the lithium acetate/PEG method described by (Gietz and Schiestl, 2007). Yeast strains expressing chromosomally Pdi1-mCherry (CY6010) or BiP-mCherry (CY6008) are described in (Young et al., 2013).

**Table 1.**
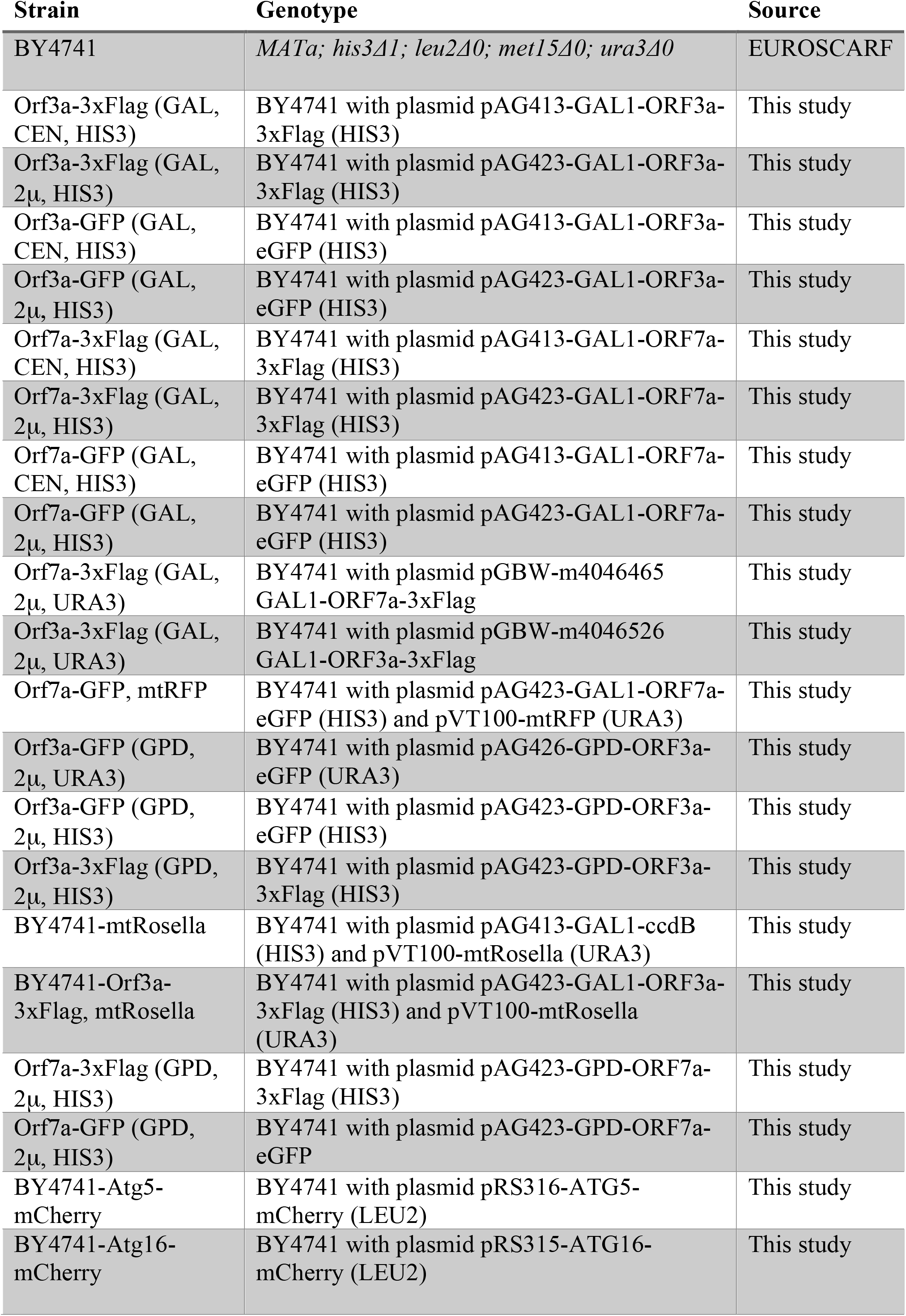
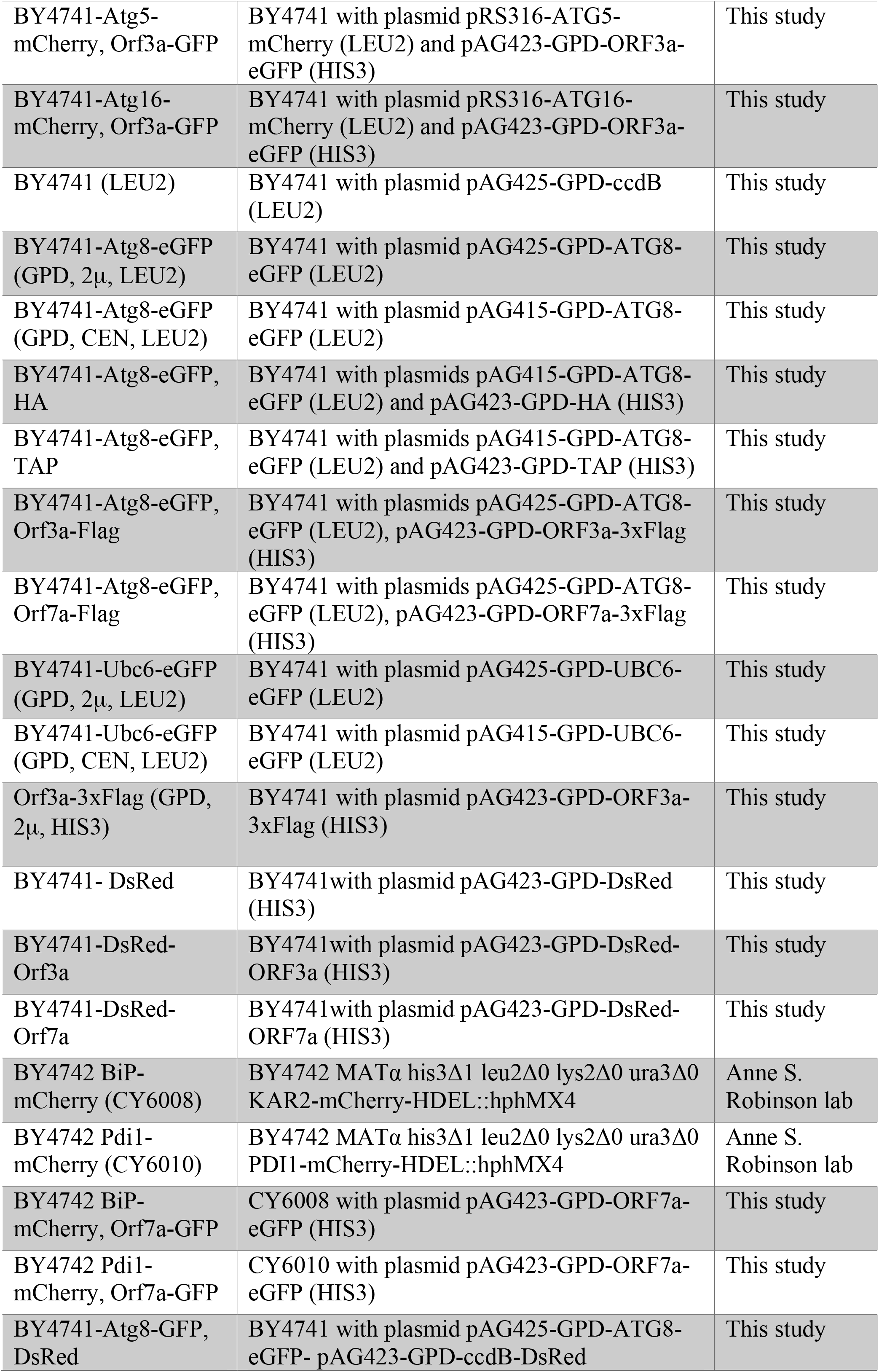
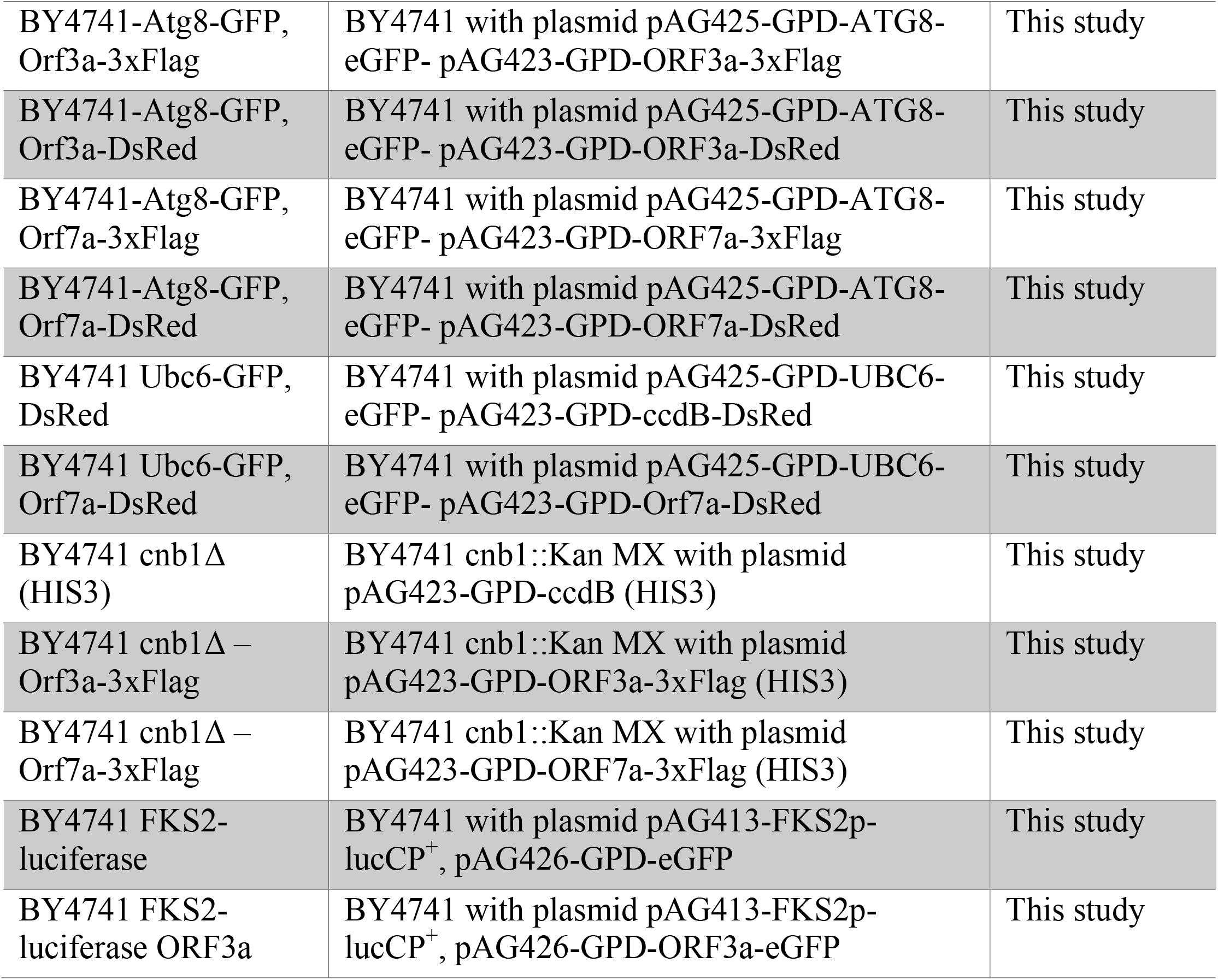
Yeast strains used in this study

### Plasmid constructions

For the galactose inducible overexpression of SARS-CoV-2 ORF3a-3xflag and ORF7a-3xflag, Addgene plasmids pGBW-m4046526 and pGBW-m4046465 were used. All other constitutive or galactose inducible ORF3a and ORF7a fusion plasmids were generated by subcloning the complete coding regions into pDONR221 and subsequent insertion by Gateway technology into the yeast destination vectors pAG413-GAL1-ccdB, pAG423-GAL1-ccdB, pAG413-GAL1-ccdB-eGFP, pAG423-GAL1-ccdB-eGFP, pAG423-GPD-ccdB-eGFP, pAG426-GPD-ccdB-eGFP, pAG423-GPD-dsRed-ccdB (Alberti et al., 2007). ATG8 and UBC6 were N-terminally tagged with eGFP by Gateway cloning of the entire ATG8 ORF or the C-terminal ER transmembrane anchor of Ubc6 into the yeast destination vectors pAG415-GPD-eGFP-ccdB or pAG425-GPD-eGFP-ccdB (Alberti et al., 2007). The live cell FKS2p-luciferase reporter was constructed by subcloning the FKS2 promoter region SacI/SmaI into the destabilized luciferase plasmid pAG413-lucCP^+^ (Rienzo et al., 2012). All plasmids were verified by DNA sequencing. Oligonucleotide primers are summarized in Table 2.

**Table 2.**
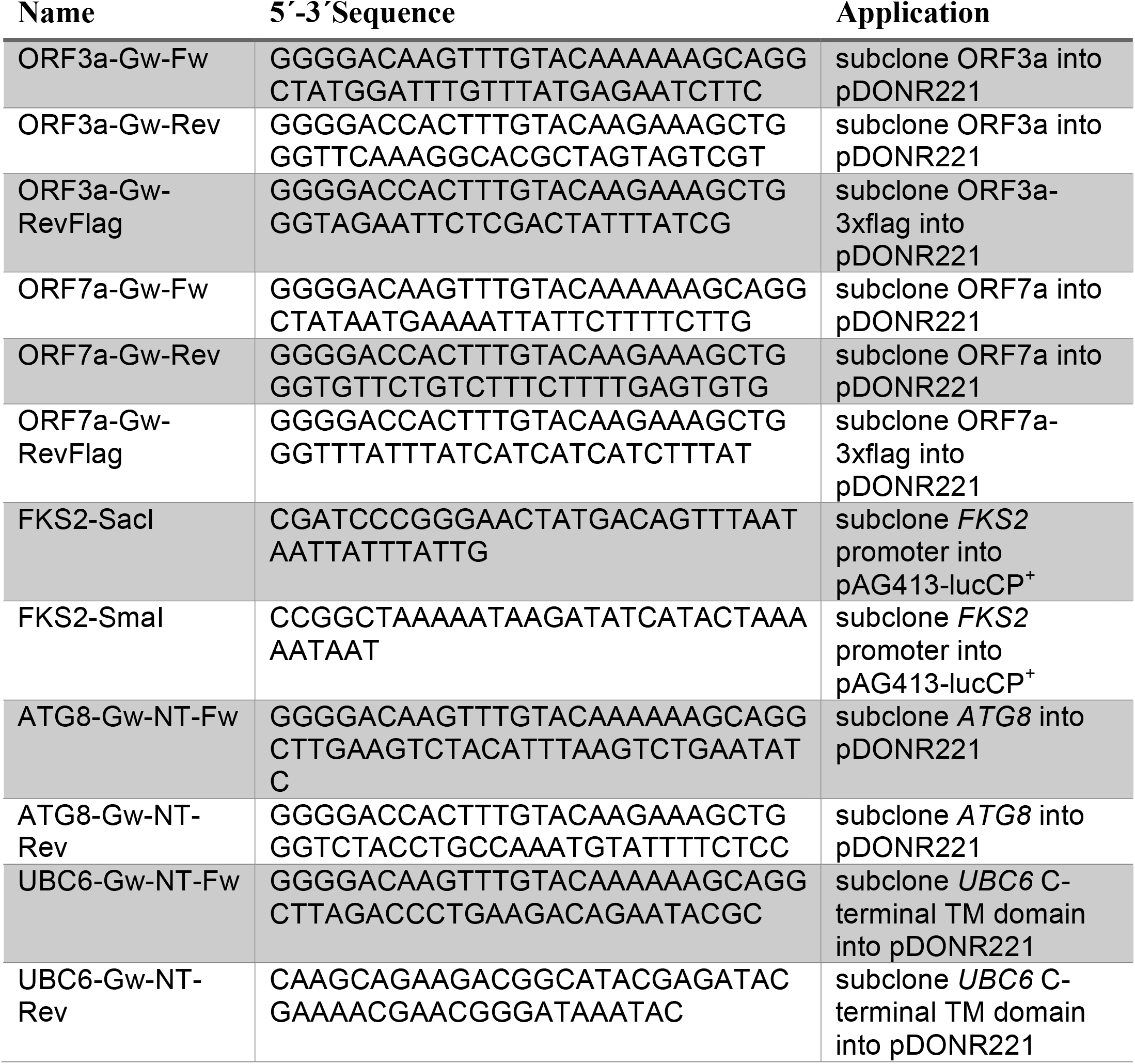
Oligonucleotide primers used in this study

### Yeast quantitative growth assays

Continuous growth curves for different yeast cultures were obtained on a Tecan Spark microplate reader. Fresh overnight SD cultures were diluted in triplicate in multiwell plates to the same starting OD. The OD_600_ was then continuously monitored for the indicated times. The growth efficiency was determined by calculating the time needed to reach the half maximal OD for each strain and condition and compared to the respective control cells or condition as indicated in the Figures.

### Immunological methods

Yeast whole cell protein extracts were prepared by the alkaline pretreatment and boiling procedure described in (Kushnirov, 2000). 2.5 OD_600_ of yeast cells were harvested and incubated with 0.1M NaOH for 5 min at room temperature. Cells were pelleted and resuspended in 50 μl of Laemmli SDS sample buffer. The samples were then boiled for 5 min at 95oC and centrifuged. 10-20μl of the supernatant was resolved on 10-15% polyacrylamide gels, transferred to PVDF membranes and probed with different antibodies. The following primary antibodies were used in this work: Rabbit anti-GFP (TP401, Amsbio), mouse anti-flag (M2, Sigma). Secondary antibodies were anti-rabbit or anti-mouse conjugated to peroxidase (Amersham Biosciences). The bands were visualized with ECL Plus and quantified with an ImageQuant LAS4000 system.

### Fluorescence microscopy

Exponentially growing yeast cells were visualized on a Leica SP8 confocal microscope with a HCXPL APO CS2 63x objective. Vacuoles were stained for 30 min with 100μM CellTracker Blue CMAC (Life Technologies) in living cells. GFP was visualized with 488 nm excitation and 509 nm emission, dsRed with 545 nm excitation and 572 nm emission, CMAC with 353 nm excitation and 466 nm emission.

### Live cell luciferase assay

Yeast strains harboring the indicated destabilized luciferase reporters were grown to exponential phase in SD supplemented with the appropriate amino acids and adjusted to pH 3.0 with 50 mM succinic acid. Cells were incubated on a roller for 60 min at 28oC with 0.5 mM luciferin (free acid, Synchem, Germany). Cell aliquots were then transferred to white 96-well plates, which contained the indicated Ca^2+^ concentrations. The light emission was immediately measured in a GloMax microplate luminometer (Promega) in triplicate and continuously recorded over the indicated time. For the ORF3a overexpressing cells, we recorded the initial 10 readings after luciferin loading of the cells.

### GFP-Atg8 autophagy assay

Yeast cells constitutively expressing GFP-Atg8 from the pAG425-GPD-eGFP-ATG8 plasmid and expressing or not ORF3a-3flag or ORF7a-3flag were grown to exponential phase. Uninduced samples were taken at time point 0, the rest of the cells were washed once with water and finally resuspended in starvation medium (SD-N) for induction of autophagy. Samples were taken at the indicated times and whole cell protein extracts prepared according to (Kushnirov, 2000). GFP-Atg8 and free GFP were visualized by western blotting and the bands quantified using the ImageJ software. The relative amount of GFP-Atg8 cleavage was quantified by determining the ratio [free GFP]/[GFP-Atg8 + free GFP].

## Results

### Overexpression of SARS-CoV-2 ORF3a and ORF7a in yeast

The small proteins encoded by the SARS-CoV-2 ORF3a (275aa; 31kD) and ORF7a (121aa; 14kD) genes are important for an efficient virus multiplication and egression from the host cell. When overexpressed separately in human cells, they decrease viability by 20-30% (Lee et al., 2021). The predicted topology indicates that both accessory proteins can insert into biological membranes via 1 (ORF7a) or 3 (ORF3a) hydrophobic transmembrane domains (Lee et al., 2021). Additionally, both proteins contain N- and C-terminal soluble domains of variable length (Figure 1A). In order to overexpress the ORF3a and ORF7a proteins in yeast, we constructed a set of centromeric or high copy expression plasmids containing C-terminal Flag-or GFP-fusion genes under the control of the inducible *GAL1* or the constitutive *TDH3* promoters (Figure 1B). We used the galactose inducible plasmids to verify the protein expression in transgenic yeast cell lines. As shown in Figure 1C, both ORF3a and ORF7a were correctly and stably overexpressed in yeast. Interestingly, while ORF7a protein content was increased by the use of high copy vectors, the maximal abundance of ORF3a was already achieved from single copy expression vectors. In any case, for the functional analysis of both small ORFs, we applied high copy constructs throughout the work. We next tested how ORF3a and ORF7a overexpression affected cell fitness in the yeast system by quantitative growth assays. We found that the induced overexpression of both, ORF3a or ORF7a inhibited cell proliferation by about 20% (Figure 1D). These results indicated that SARS-CoV-2 ORF3a and ORF7a proteins are biologically active when overexpressed in yeast cells.

**Figure 1:**
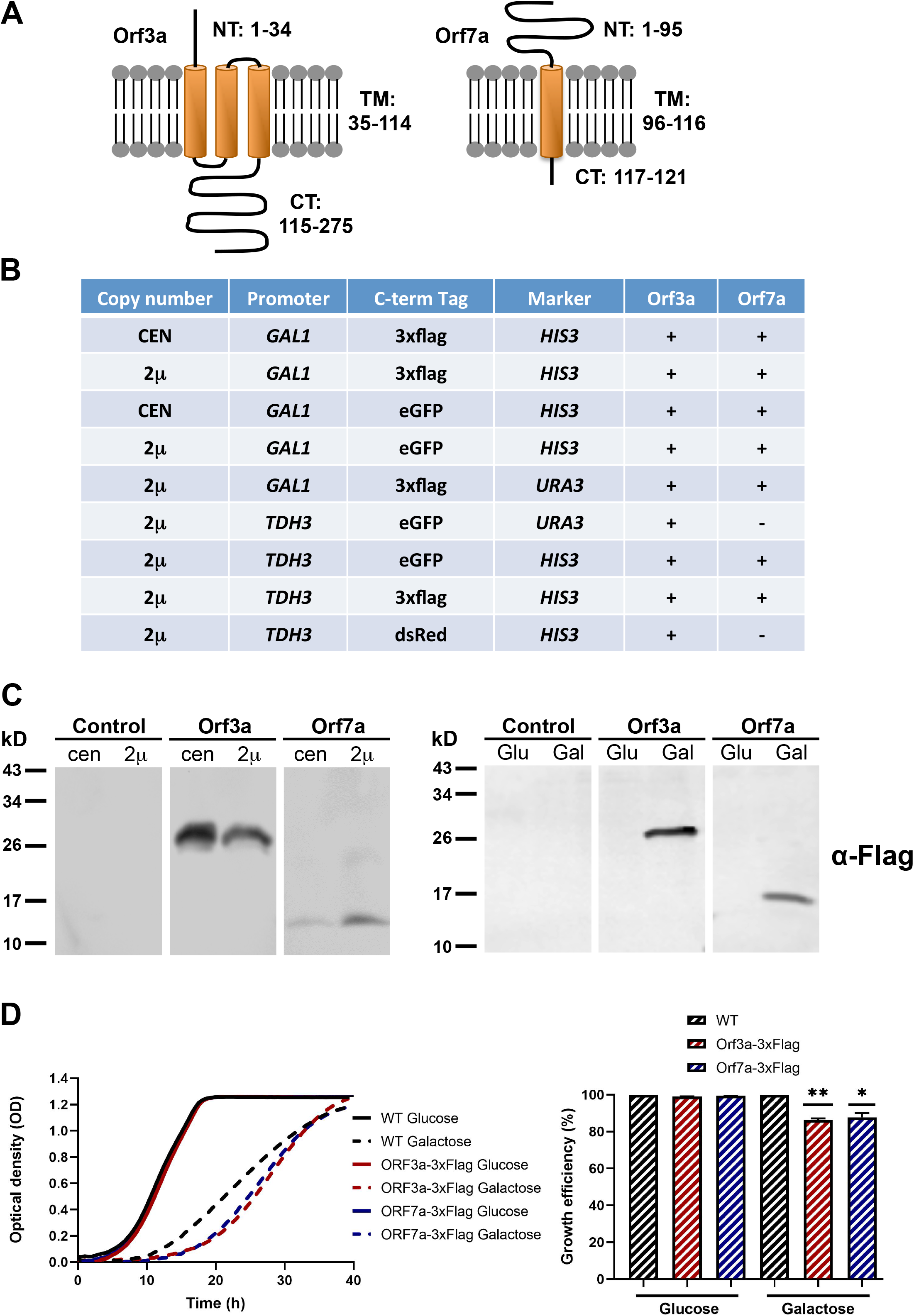
Overexpression of the SARS-CoV-2 small accessory proteins ORF3a and ORF7a in yeast. (A) Predicted topolgy of membrane inserted SARS-CoV ORF3a and ORF7a. NT = N-terminal soluble domain; TM = Trans membrane hydrophobic domain; CT = C-terminal soluble domain. Numbers indicate the amino acids comprising each predicted domain. (B) Heterologous yeast expression systems for the overexpression of differentially C-terminally tagged ORF3a and ORF7a proteins created in this work. CEN = centromeric plasmid; 2μ = high copy plasmid; GAL1 = galactose inducible GAL1 promoter; TDH3 = constitutively active TDH3 promoter. (C) Immunological detection of constitutively (left panel) and galactose-induced (right panel) expression of SARS-CoV-2 ORF3a-flag and ORF7a-flag fusion proteins in yeast whole cell extracts. (D) Growth inhibition of SARS-CoV-2 ORF3a and ORF7a overexpression in yeast wild type cells. High copy, galactose-inducible expression plasmids were used. Left panel: Continuous growth curve determination; right panel: Representation of the relative growth efficiency. Growth of the empty plasmid control cells was set to 100% for glucose and galactose. Three independent biological samples were analyzed. Data are mean +/-SD. * p < 0.05; ** p < 0.01 (unpaired Student’s t-test).

### Intracellular localization of SARS-CoV-2 ORF3a and ORF7a in yeast

We visualized the intracellular distribution of constitutively overexpressed ORF3a-GFP and ORF7a-GFP fusion proteins by confocal fluorescence microscopy. We found that both SARS-CoV-2 small ORF proteins were targeted to distinct cellular sub-compartments. ORF3a localized in all cells to the vacuolar membrane (Figure 2), which in many cases showed a fragmented architecture. Additionally, ORF3a was detected in intracellular deposits, whose number varied considerably from cell to cell. We did not find ORF3a-GFP inside vacuoles, which indicated that ORF3a specifically inserted in the vacuolar membrane to exert its biological function.

**Figure 2:**
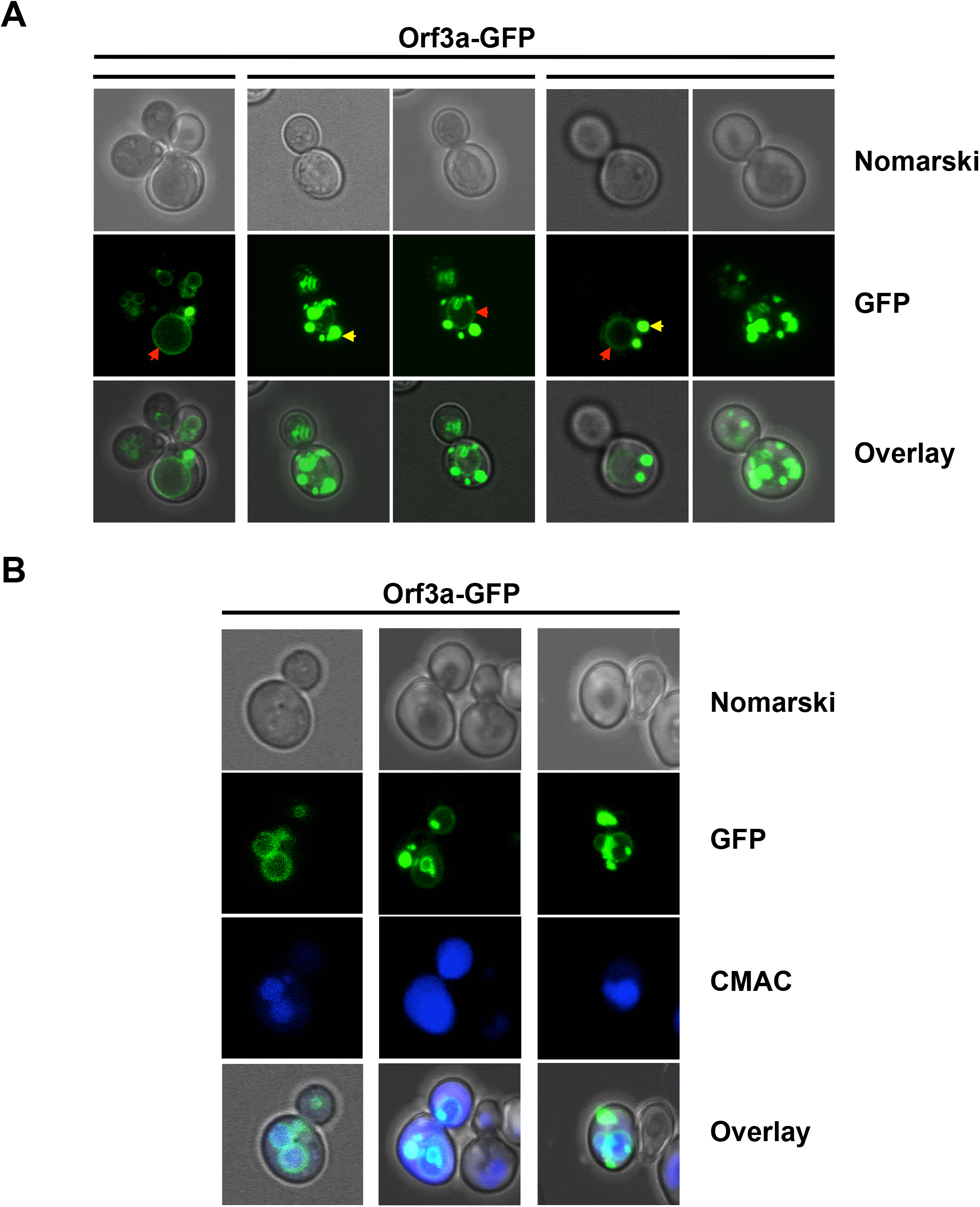
SARS-CoV-2 ORF3a localizes to the vacuolar membrane in yeast cells. ORF3a-GFP was constitutively overexpressed in yeast wild type cells from high copy plasmids and visualized by confocal fluorescence microscopy. (A) Representative examples of ORF3a-GFP intracellular distribution. Different z-stacks are represented as indicated. Red arrows mark an apparent vacuolar envelope localization, yellow arrows mark a localization to round deposits for ORF3a. (B) ORF3a-GFP colocalization with the vacuolar membrane.

ORF7a-GFP was visualized in yeast cells in an intracellular diffuse/punctate structure, which resembled the endoplasmic reticulum (ER). Therefore we created two-colored reporter yeast strains, which combined the overexpression of ORF7a fused to GFP or dsRed with red and green ER-marker proteins. To this end we applied an N-terminal GFP-fusion with the Ubc6 ER-specific membrane anchor and a C-terminal mCherry fusion of the ER resident Kar2 (BiP) chaperone. In both combinations, we found that the ORF7a and ER-marker signals largely coincided (Figure 3). These results indicated that ORF7a targets predominantly the ER in yeast cells, most likely by insertion into the ER membrane via its C-terminal TM domain. We next addressed the question whether overexpressed SARS-CoV-2 ORF7a caused ER damage. We used different doses of the ER inhibitor tunicamycin in control and constitutively ORF7a expressing yeast cells (Figure 3B). In this assay, an increased susceptibility to tunicamycin in the presence of ORF7a could identify an interference of this small ORF with general ER functions. However, we only noticed a moderate growth inhibition caused by ORF7a expression already upon control conditions. The inhibitory effect of tunicamycin was not further exacerbated in the overexpressor cell lines (Figure 3B). We conclude that SARS-CoV-2 ORF7a inserts specifically into the yeast ER membrane system without causing a detectable malfunction at this organelle.

**Figure 3:**
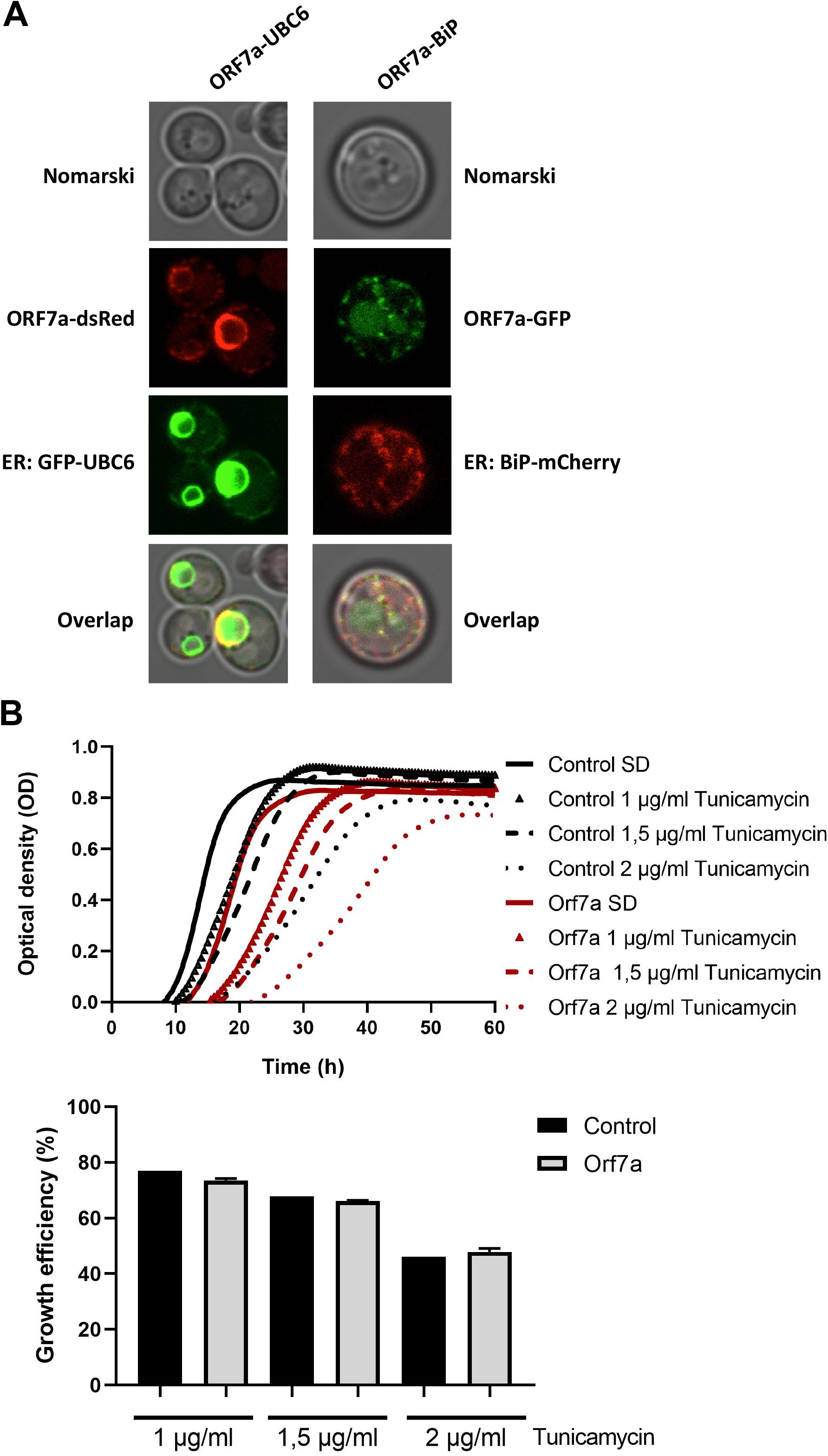
SARS-CoV-2 ORF7a localizes to the ER in yeast cells. (A) ORF7a-GFP or ORF-7a-dsRed was constitutively overexpressed in yeast wild type cells from high copy plasmids and visualized by confocal fluorescence microscopy. GFP-Ubc6 or BiP-mCherry were co-expressed as green or red ER markers. (B) ER localization of SARS-CoV-2 ORF7a does not confer a general ER dysfunction. Constitutively ORF7a overexpressing yeast cells were treated or not with the indicated concentrations of ER inhibitor tunicamycin. Upper panel: Continuous growth curves derived from 3 biological replicates. Lower panel: Representation of the relative growth efficiency. Growth of the empty plasmid control cells was set to 100% for SD medium without tunicamycin. Three independent biological samples were analyzed. Data are mean +/-SD.

### SARS-CoV-2 ORF3a and ORF7a interfere with the autophagic flux via different mechanisms

Autophagy is a major host cell function, which is often modulated by viral accessory proteins. Having localized SARS-CoV-2 ORF7a at the ER network, which is a principal localization of emerging autophagosomes (Hollenstein and Kraft, 2020), and ORF3a at the vacuolar membrane, which is the final destination of mature autophagosomes in yeast, we investigated, whether the overexpression of both virus factors influenced the process of autophagy in yeast cells. We first looked at the steady state appearance of autophagosomal structures by the visualization of GFP-Atg8 containing vesicles in live yeast cells. Favorable growth conditions in the presence of abundant fermentable sugars repress autophagy in yeast (Adachi et al., 2017). Therefore, in proliferating yeast populations, cells normally do not contain detectable autophagosomal vesicles (Figure 4A). However, when SARS-CoV-2 small ORFs 3a or 7a were overexpressed, in the majority of the cells one or two GFP-Atg8 associated vesicles could be visualized. Thus, both SARS-CoV-2 accessory proteins increased the number of cytosolic autophagosomal structures in yeast, which in principal could be the consequence of either an enhanced formation or a reduced processing of autophagosomes.

**Figure 4:**
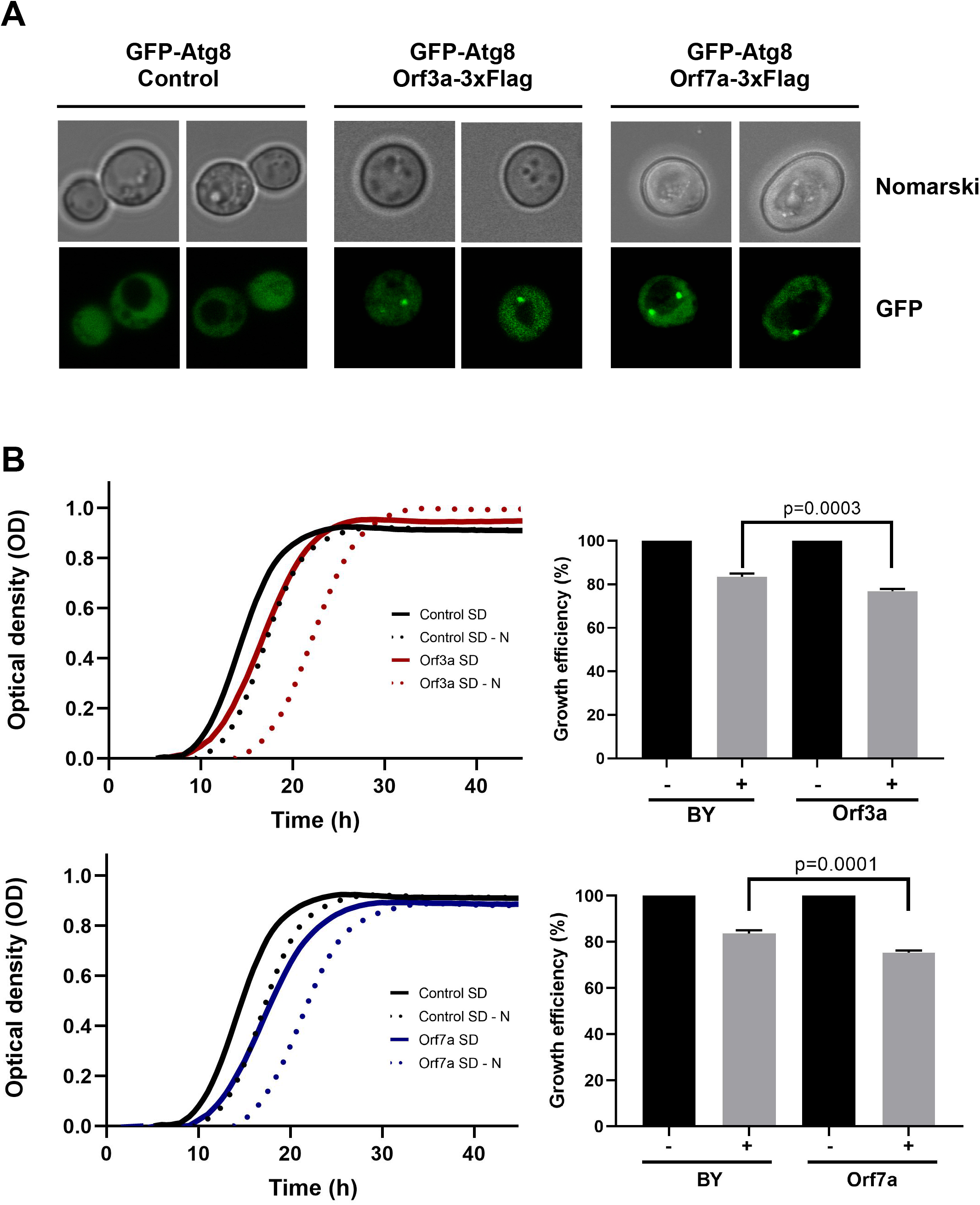
Overexpression of SARS-CoV-2 ORF3a and ORF7a leads to accumulation of Atg8-positive autophagosomes and sensitivity to starvation. (A) The formation of autophagosomal structures was visualized by the expression of GFP-tagged Atg8 in yeast wild type cells in the absence (empty vector control) or the presence of constitutively overexpressed Orf3a-flag or Orf7a-flag as indicated. Cells were grown to exponential phase in SD medium. Representative micrographs are shown for each genetic background. (B) SARS-CoV-2 ORF3a (upper panel) and ORF7a (lower panel) were constitutively overexpressed as flag fusion proteins in yeast wild type cells and the growth continuously monitored in SD medium. Cells were either directly diluted from SD medium (-starvation) or after three days in SD-N medium (+ starvation). Control cells contain the empty plasmid. Right panels: Representation of the relative growth efficiency. Growth of the non starved cells was set to 100%. Three independent biological samples were analyzed. Data are mean +/-SD. P-values indicate significant differences between control and overexpressor cells according to the unpaired Student’s t-test.

Autophagic clearance of cellular components and waste becomes essential in yeast upon starvation conditions. We therefore tested whether the overexpression of ORF3a or ORF7a limited the survival of yeast cells during nutrient starvation conditions by quantifying the growth efficiency after a prolonged incubation upon nitrogen starvation. As shown in Figure 4B, both the accumulation of ORF3a and ORF7a, had a significant negative effect on the fitness of yeast during nitrogen starvation. These results suggested that both viral accessory proteins manipulated the autophagic process in a physiological manner. In order to quantify the effect of ORF3a and ORF7a on autophagic processing, we immunologically measured the quantity of digested GFP-Atg8 protein in the vacuole by the appearance of free GFP (Araki et al., 2017). As shown in Figure 5, induction of autophagic GFP-Atg8 cleavage by starvation is readily quantified in yeast control cells. The overexpression of ORF3a significantly decreased the GFP-Atg8 autophagic processing, indicating that ORF3a, once localized at the yeast vacuolar membrane, impairs here the fusion and final processing of autophagosomes. In the case of ORF7a overexpression, we observed a significant stimulation of basal and induced autophagic GFP-Atg8 cleavage (Figure 5). These findings suggest that both SARS-CoV-2 small ORFs efficiently interfere with yeast autophagic flux, ORF7a by stimulating autophagosomal formation at the ER and ORF3a by decreasing vacuolar processing of autophagosomes.

**Figure 5:**
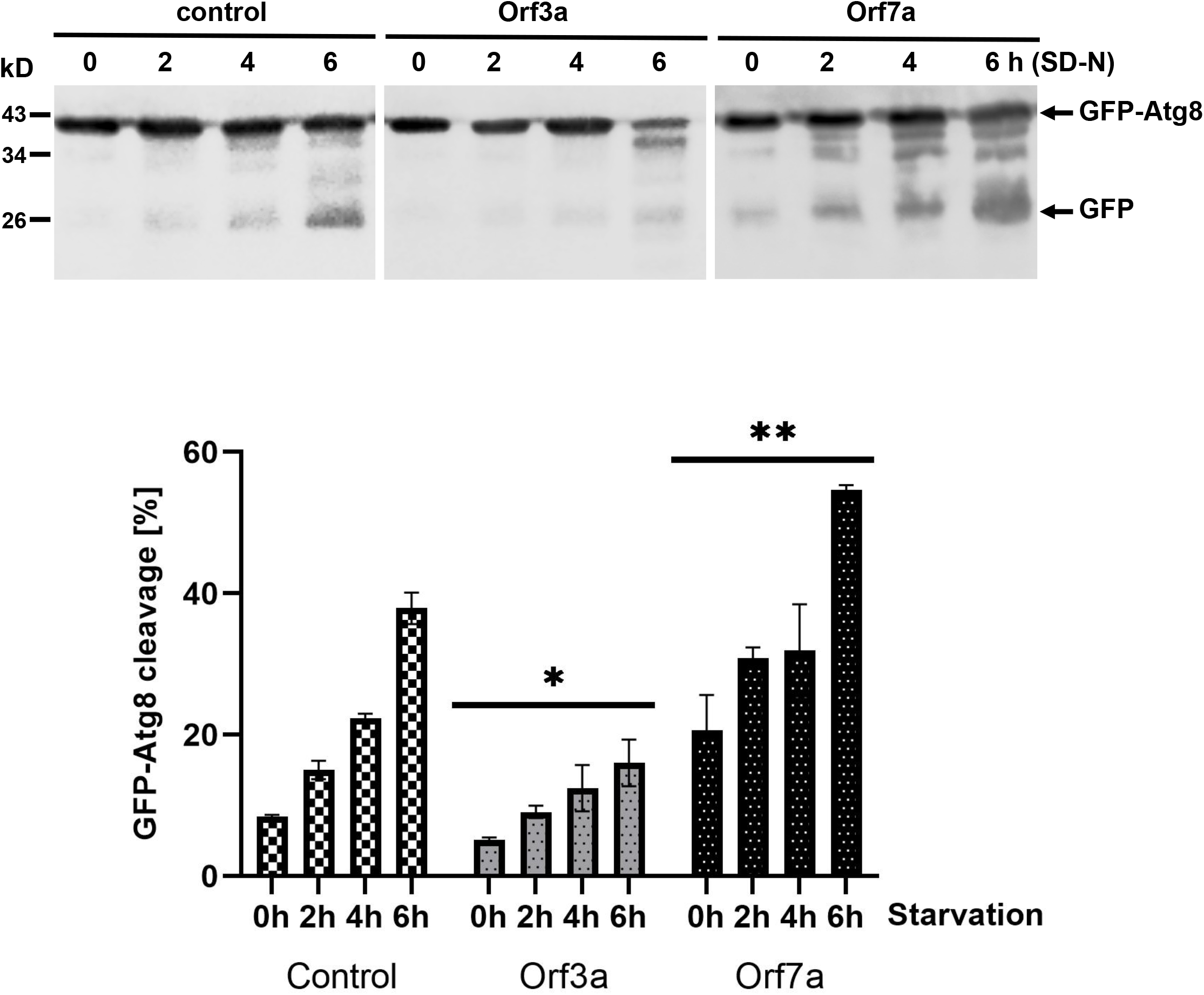
Effects of SARS-CoV-2 ORF3a and ORF7a overexpression on the autophagic digestion of GFP-Atg8. GFP-tagged Atg8 was constitutively expressed in yeast cells in the presence or absence (control) of ORF3a or ORF7a. Upper panel: Immunological detection of full length GFP-Atg8 or proteolytically cleaved GFP before and during starvation in SD-N medium. Lower panel: Representation of the relative GFP-Atg8 cleavage. Three independent biological samples were analyzed. Data are mean +/-SD. * p < 0.05; ** p < 0.01 (unpaired Student’
ss t-test) for significant differences between control and overexpressor cell lines.

### SARS-CoV-2 ORF3a alters Ca^2+^ homeostasis in yeast

The purification and functional reconstitution of the SARS-CoV-2 ORF3a protein has recently suggested that this viral protein could act as a non-specific, Ca^2+^ permeable, ion channel in host cells (Kern et al., 2021). Our results localize ORF3a at the vacuolar membrane in transfected yeast cells. Since the vacuoles accumulate >90% of the cellular calcium and thus are the major Ca^2+^ storage compounds in yeast (Cunningham and Fink, 1994), we set out to investigate whether SARS-CoV-2 ORF3a modulated the Ca^2+^ flux at vacuoles *in vivo*. First, we determined the tolerance of yeast cells to different externally added Ca^2+^ doses in the presence or absence of overexpressed ORF3a by quantitative growth assays. As depicted in Figure 6A, the constitutive overexpression of SARS-CoV-2 ORF3a in yeast wild type cells caused a mild growth delay upon normal growth conditions. Interestingly, we observed a significant improvement of growth upon high Ca^2+^ concentrations in the presence of overexpressed ORF3a. The evolutionarily conserved calmodulin-calcineurin signaling pathway is responsible in yeast to adjust intracellular Ca^2+^ homeostasis (Thewes, 2014; Park et al., 2019). Its central protein phosphatase calcineurin is activated by elevated cytosolic Ca^2+^ concentrations and orchestrates cellular stress responses, which include stimulated Ca^2+^ transport in order to adapt to Ca^2+^ overload. We hypothesized that Ca^2+^ tolerance in the ORF3a overexpressing yeast cells could be caused by activation of the calmodulin-calcineurin signal transduction pathway. To test this hyptothesis, we overexpressed SARS-CoV-2 ORF3a in a *cnb1* mutant lacking the function of the calcineurin B subunit of the calcineurin complex. We found that in the absence of calcineurin signaling the overexpression of ORF3a inhibited proliferation more strongly already upon normal Ca^2+^ conditions, which was further exacerbated by increasing external Ca^2+^ addition (Figure 6B). These data suggested that SARS-CoV-2 ORF3a caused Ca^2+^ leakage from yeast vacuoles, which stimulated an adaptive response via the calmodulin-calcineurin pathway (Figure 6C). To further test this model, we sought to measure the expression levels of Ca^2+^-calmodulin-calcineurin regulated target genes in the presence or absence of ORF3a. One of the most sensitively Ca^2+^ up-regulated target genes is the *FKS2* gene encoding a catalytic subunit of 1,3-beta-glucan synthase at the cell membrane (Stathopoulos and Cyert, 1997). In order to create a Ca^2+^ sensitive live cell reporter, we fused the *FKS2* upstream control region with a destabilized version of firefly luciferase. The resulting FKS2-luciferase reporter was indeed efficient to detect Ca^2+^ stimulated gene expression in live yeast cells (Figure 6D). We finally overexpressed SARS-CoV-2 ORF3a in yeast wild type cells harboring the FKS2-luciferase reporter. As shown in Figure 6E, a significantly elevated luciferase activity was consistently found in different ORF3a overexpressing yeast cell lines, indicating that SARS-CoV-2 ORF3a activated Ca^2+^ dependent gene expression most likely by promoting Ca^2+^ leakage from the vacuole.

**Figure 6:**
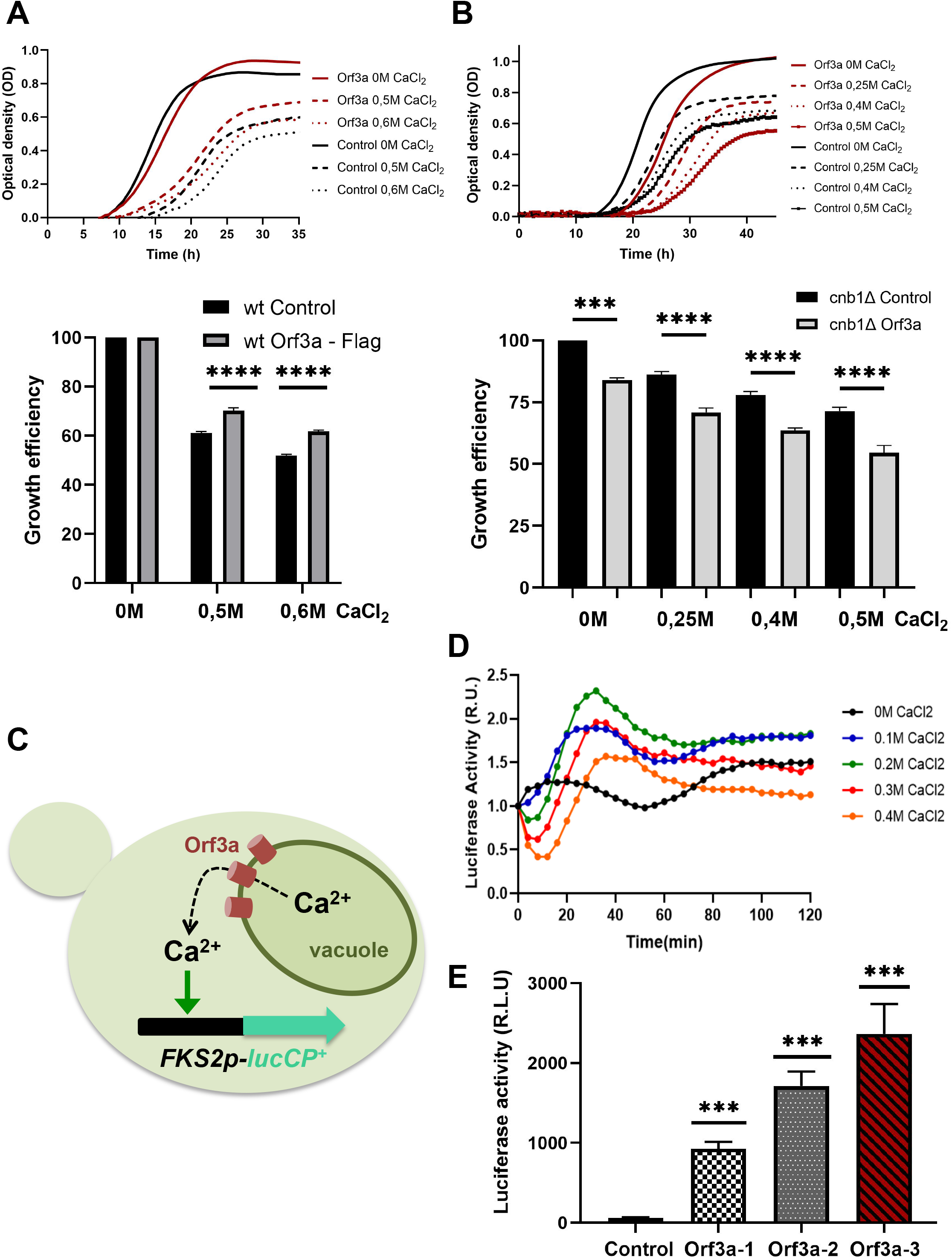
SARS-CoV-2 ORF3a modulates intracellular Ca^2+^ homeostasis in yeast. (A) ORF3a confers Ca^2+^ tolerance in yeast wild type cells. Upper panel: Growth of constitutively ORF3a overexpressing or control cells in SD medium with the indicated Ca^2+^ concentrations. Lower panel: Representation of the relative growth efficiency. Growth of the control cells without addition of Ca^2+^ was set to 100%. (B) ORF3a confers Ca^2+^ sensitivity in yeast *cnb1* mutants defective for calcineurin signaling. Experimental procedure same as in (A). Three independent biological samples were analyzed. Data are mean +/-SD. (C) Experimental model connecting ORF3a mediated Ca^2+^ efflux and its detection by the FKS2-luciferase live cell reporter. (D) The FKS2-lucCP^+^ reporter is activated by Ca^2+^ in a dose-dependent manner. Yeast wild type cells harboring the FKS2-luciferase reporter plasmid were grown in SD medium and the indicated Ca^2+^ doses added at time point 0. The luciferase activity was determined continuously by the light emission from live cell populations. Data are mean values from three biological replicates. (E) Constitutively overexpressed ORF3a activates the Ca^2+^ sensitive FKS2-luciferase reporter. Three different ORF3a overexpressing yeast cell lines were compared to control cells. Three biological replicates were analyzed. Data are mean +/-SD. (A, B, E) p-values indicate significant differences between control and overexpressor cells according to the unpaired Student’s t-test; *** p < 0.001; **** p < 0.0001.

## Discussion

Budding yeast is an attractive model system to study virus host cell interactions by the heterologous expression of virulence factors (Zhao, 2017). This approach, however, can only discover meaningful insights in cases where the viral proteins target cellular processes, which are evolutionarily conserved enough to simulate the virus-host interaction in a simple yeast cell. For example, several viruses manipulate the host cell cycle or programmed cell death pathways in a way to maximize cellular resources for their own reproduction (Hay and Kannourakis, 2002; Swanton and Jones, 2001). Both cell cycle and apoptosis, are important targets for virulence factors, which have been successfully studied in budding and fission yeast models for human viruses such as HIV-1, human papilloma virus or Zika virus (Zhao et al., 1996; Li et al., 2017; Macreadie et al., 1995; Fournier et al., 1999). Here we show that SARS-CoV-2 accessory proteins ORF3a and ORF7a, when expressed in budding yeast cells, are stable, correctly localized to discrete membrane compartments and display defined biological functions in autophagy and calcium homeostasis. Autophagy is evolutionarily highly conserved from fungi to humans, and provides virus-infected cells with an innate defense mechanism by selectively clearing virions or individual viral proteins or genomes via lysosomal degradation (Deretic et al., 2013). Furthermore, in the specific case of β-coronavirus propagation, the virus hijacks and exploits the lysosomal exocytosis pathway for its own egression and spreading from the host cell (Ghosh et al., 2020). Thus, efficient SARS-CoV-2 propagation depends on the simultaneous stimulation of autophagosome formation and inhibition of autophagosome maturation and lysosomal degradation (Chen and Zhang, 2022). We show that the SARS-CoV-2 accessory proteins ORF3a and ORF7a play complementary roles to achieve this dysfunction of the autophagy process. ORF3a targets specifically the vacuolar membrane in yeast, from where it causes the accumulation of autophagosomal structures by impairing their final degradation at vacuoles. This function is analogous to what has been found in human cells, where ORF3a localizes to the late endosome/lysosomes, the functional homologue of yeast vacuoles, and increases the number of cytosolic autophagosomal vesicles (Miao et al., 2021). This is due to the capacity of SARS-CoV-2 ORF3a to interact with the HOPS component Vps39 and thereby block the fusion of lysosomes with autophagosomes via the Stx17/Snap29/Vamp8 SNARE complex (Miao et al., 2021; Zhang et al., 2021). We assume that this specific function also occurs in yeast cells because the vacuolar HOPS complex is highly conserved in fungi (Balderhaar and Ungermann, 2013). Interestingly, the interaction of ORF3a with the HOPS Vps39 subunit seems to represent a recent gain of function of this accessory protein, because it is absent in the SARS-CoV-1 counterpart (Miao et al., 2021; Chen et al., 2021) and thus might not be the ancestral function of Coronavirus ORF3a proteins. Additionally, SARS-CoV ORF3a proteins form a sophisticated structure at membranes, where a homodimeric complex with a total of 6 transmembrane domains has a predicted non-selective Ca^2+^ permeable cation channel activity (Kern et al., 2021). This is a complexity that would probably not be necessary if the sole function of ORF3a was to present an interfering domain for Vps39 interaction at lysosomes/vacuoles. Here we present several evidences that SARS-CoV-2 ORF3a disturbs Ca^2+^ homeostasis in yeast, because its overexpression causes Ca^2+^ resistance dependent on calcineurin/calmodulin signaling and activates a cytosolic Ca^2+^ sensitive reporter. Given the selective localization of ORF3a at the vacuolar membrane, this suggests that SARS-CoV-2 ORF3a causes Ca^2+^ leakage from the vacuole, which stimulates Ca^2+^ homeostatic adaptation through calcineurin. Therefore, SARS-CoV-2 ORF3a might indeed be a bona fide viroporin with Ca^2+^ channel activity *in vivo*, which yet has to be confirmed in human cells. Several viroporins from other viruses have been shown to manipulate Ca^2+^ homeostasis for the benefit of their own multiplication (Nieva et al., 2012). Specificallly, it is well known for some viroporins that their Ca^2+^ release function is aimed at the stimulation of autophagy via activation of Ca^2+^/calmodulin and the AMP-activated protein kinase AMPK (Crawford et al., 2012; Crawford and Estes, 2013; Tokumitsu and Sakagami, 2022). An exciting hypothesis could be that SARS-CoV-2 ORF3a might have evolved as a dual function autophagy modulator, (i) stimulating autophagy indirectly via Ca^2+^ release from vacuoles/lysosomes and (ii) inhibiting directly the processing of autophagosomes at vacuoles/lysosomes by interfering with SNARE complex binding. Additionally, the release of lysosomal calcium might be the trigger SARS-CoV-2 ORF3a uses to induce the previously reported entry into apoptotic cell death programs (La Rovere et al., 2016; Ren et al., 2020).

SARS-CoV-2 ORF7a contributes yet other pro-autophagic functions, which in combination with the ORF3a functions might lead to a multi-functional and optimized induction of incomplete autophagy in the host cell. SARS-CoV-2 ORF7a is targeted to yeast ER membranes by just one transmembrane domain. Specificity of SARS-CoV-2 ORF7a localization might be conferred by an N-terminal 15aa cleavable signal peptide and a C-terminal ER retention sequence KRKTE originally identified in SARS-CoV-1 (Fielding et al., 2004). The ORF7a localization reported here in yeast host cells is in agreement with the finding that SARS-CoV-2 ORF7a is directed to the ER and Golgi apparatus in human cells (Lee et al., 2021; Zhang et al., 2020). Here we report a stimulating function of SARS-CoV-2 ORF7a on the formation of Atg8 containing autophagosomes, which are probably derived from the ER. The induction of double membrane-coated vesicles (DMVs) from the ER is an important mechanism for coronaviruses to provide host cell vehicles for their genome replication and protection from immune defenses (Reggiori et al., 2010; Wolff et al., 2020). Thus, SARS-CoV-2 ORF7a could be the accessory protein in charge of providing ER derived DMVs for intracellular trafficking of the virus. ORF3a mediated inhibition of vesicle fusion with vacuoles/lysosomes on the other hand would enhance the possibilities of the virus to egress and spread via the induction of lysosomal exocytosis (Chen et al., 2021; Solvik et al., 2022). Our work shows that SARS-CoV-2 accessory proteins can be functionally studied in yeast cells to gain insights into their biological functions, in the case reported here, by manipulating autophagic processes between different membrane compartments with the participation of Ca^2+^ homeostasis.

## Acknowledgments

The authors thank Anne S. Robinson for the kind gift of BiP-mCherry and Pdi1-mCherry expressing yeast strains and Xi Zhou for the kind gift of SARS-CoV-2 ORF3a expression plasmid pHAGE-ORF3a-flag.

## Bibliography

Abdoli, A., Alirezaei, M., Mehrbod, P., and Forouzanfar, F. (2018). Autophagy: The multi-purpose bridge in viral infections and host cells. Rev Med Virol 28, e1973. doi:10.1002/rmv.1973.

Adachi, A., Koizumi, M., and Ohsumi, Y. (2017). Autophagy induction under carbon starvation conditions is negatively regulated by carbon catabolite repression. J. Biol. Chem. 292, 19905–19918. doi:10.1074/jbc.M117.817510.

Alberti, S., Gitler, A. D., and Lindquist, S. (2007). A suite of Gateway cloning vectors for high-throughput genetic analysis in Saccharomyces cerevisiae. Yeast 24, 913–919. doi:10.1002/yea.1502.

Araki, Y., Kira, S., and Noda, T. (2017). Quantitative Assay of Macroautophagy Using Pho8△60 Assay and GFP-Cleavage Assay in Yeast. Meth. Enzymol. 588, 307– 321. doi:10.1016/bs.mie.2016.10.027.

Balderhaar, H. J. kleine, and Ungermann, C. (2013). CORVET and HOPS tethering complexes - coordinators of endosome and lysosome fusion. J. Cell Sci. 126, 1307–1316. doi:10.1242/jcs.107805.

Chen, D., and Zhang, H. (2022). Autophagy in severe acute respiratory syndrome coronavirus 2 infection. Curr. Opin. Physiol. 29, 100596. doi:10.1016/j.cophys.2022.100596.

Chen, D., Zheng, Q., Sun, L., Ji, M., Li, Y., Deng, H., and Zhang, H. (2021). ORF3a of SARS-CoV-2 promotes lysosomal exocytosis-mediated viral egress. Dev. Cell 56, 3250–3263.e5. doi:10.1016/j.devcel.2021.10.006.

Crawford, S. E., and Estes, M. K. (2013). Viroporin-mediated calcium-activated autophagy. Autophagy 9, 797–798. doi:10.4161/auto.23959.

Crawford, S. E., Hyser, J. M., Utama, B., and Estes, M. K. (2012). Autophagy hijacked through viroporin-activated calcium/calmodulin-dependent kinase kinase-β signaling is required for rotavirus replication. Proc. Natl. Acad. Sci. USA 109, E3405–13. doi:10.1073/pnas.1216539109.

Cunningham, K. W., and Fink, G. R. (1994). Ca2+ transport in Saccharomyces cerevisiae. J. Exp. Biol. 196, 157–166. doi:10.1242/jeb.196.1.157.

Deretic, V., Saitoh, T., and Akira, S. (2013). Autophagy in infection, inflammation and immunity. Nat. Rev. Immunol. 13, 722–737. doi:10.1038/nri3532.

Fielding, B. C., Tan, Y.-J., Shuo, S., Tan, T. H. P., Ooi, E.-E., Lim, S. G., Hong, W., and Goh, P.-Y. (2004). Characterization of a unique group-specific protein (U122) of the severe acute respiratory syndrome coronavirus. J. Virol. 78, 7311–7318. doi:10.1128/JVI.78.14.7311-7318.2004.

Fournier, N., Raj, K., Saudan, P., Utzig, S., Sahli, R., Simanis, V., and Beard, P. (1999). Expression of human papillomavirus 16 E2 protein in Schizosaccharomyces pombe delays the initiation of mitosis. Oncogene 18, 4015–4021. doi:10.1038/sj.onc.1202775.

Ghosh, S., Dellibovi-Ragheb, T. A., Kerviel, A., Pak, E., Qiu, Q., Fisher, M., Takvorian, P. M., Bleck, C., Hsu, V. W., Fehr, A. R., et al. (2020). β-Coronaviruses Use Lysosomes for Egress Instead of the Biosynthetic Secretory Pathway. Cell 183, 1520–1535.e14. doi:10.1016/j.cell.2020.10.039.

Gietz, R. D., and Schiestl, R. H. (2007). High-efficiency yeast transformation using the LiAc/SS carrier DNA/PEG method. Nat. Protoc. 2, 31–34. doi:10.1038/nprot.2007.13.

Gordon, D. E., Jang, G. M., Bouhaddou, M., Xu, J., Obernier, K., White, K. M., O’Meara, M. J., Rezelj, V. V., Guo, J. Z., Swaney, D. L., et al. (2020). A SARS-CoV-2 protein interaction map reveals targets for drug repurposing. Nature 583, 459–468. doi:10.1038/s41586-020-2286-9.

Hay, S., and Kannourakis, G. (2002). A time to kill: viral manipulation of the cell death program. J. Gen. Virol. 83, 1547–1564. doi:10.1099/0022-1317-83-7-1547.

Hayn, M., Hirschenberger, M., Koepke, L., Nchioua, R., Straub, J. H., Klute, S., Hunszinger, V., Zech, F., Prelli Bozzo, C., Aftab, W., et al. (2021). Systematic functional analysis of SARS-CoV-2 proteins uncovers viral innate immune antagonists and remaining vulnerabilities. Cell Rep. 35, 109126. doi:10.1016/j.celrep.2021.109126.

Hollenstein, D. M., and Kraft, C. (2020). Autophagosomes are formed at a distinct cellular structure. Curr. Opin. Cell Biol. 65, 50–57. doi:10.1016/j.ceb.2020.02.012.

Hou, P., Wang, X., Wang, H., Wang, T., Yu, Z., Xu, C., Zhao, Y., Wang, W., Zhao, Y., Chu, F., et al. (2022). The ORF7a protein of SARS-CoV-2 initiates autophagy and limits autophagosome-lysosome fusion via degradation of SNAP29 to promote virus replication. Autophagy. doi:10.1080/15548627.2022.2084686.

Huang, C., Wang, Y., Li, X., Ren, L., Zhao, J., Hu, Y., Zhang, L., Fan, G., Xu, J., Gu, X., et al. (2020). Clinical features of patients infected with 2019 novel coronavirus in Wuhan, China. Lancet 395, 497–506. doi:10.1016/S0140-6736(20)30183-5.

Jordan, T. X., and Randall, G. (2012). Manipulation or capitulation: virus interactions with autophagy. Microbes Infect. 14, 126–139. doi:10.1016/j.micinf.2011.09.007.

Kern, D. M., Sorum, B., Mali, S. S., Hoel, C. M., Sridharan, S., Remis, J. P., Toso, D. B., Kotecha, A., Bautista, D. M., and Brohawn, S. G. (2021). Cryo-EM structure of SARS-CoV-2 ORF3a in lipid nanodiscs. Nat. Struct. Mol. Biol. 28, 573–582. doi:10.1038/s41594-021-00619-0.

Kobayashi, S., Orba, Y., Yamaguchi, H., Takahashi, K., Sasaki, M., Hasebe, R., Kimura, T., and Sawa, H. (2014). Autophagy inhibits viral genome replication and gene expression stages in West Nile virus infection. Virus Res. 191, 83–91. doi:10.1016/j.virusres.2014.07.016.

Koepke, L., Hirschenberger, M., Hayn, M., Kirchhoff, F., and Sparrer, K. M. (2021). Manipulation of autophagy by SARS-CoV-2 proteins. Autophagy 17, 2659– 2661. doi:10.1080/15548627.2021.1953847.

Kushnirov, V. V. (2000). Rapid and reliable protein extraction from yeast. Yeast 16, 857–860. doi:10.1002/1097-0061(20000630)16:9<857::AID-YEA561>3.0.CO;2-B.

La Rovere, R. M. L., Roest, G., Bultynck, G., and Parys, J. B. (2016). Intracellular Ca(2+) signaling and Ca(2+) microdomains in the control of cell survival, apoptosis and autophagy. Cell Calcium 60, 74–87. doi:10.1016/j.ceca.2016.04.005.

Lee, J.-G., Huang, W., Lee, H., van de Leemput, J., Kane, M. A., and Han, Z. (2021). Characterization of SARS-CoV-2 proteins reveals Orf6 pathogenicity, subcellular localization, host interactions and attenuation by Selinexor. Cell Biosci. 11, 58. doi:10.1186/s13578-021-00568-7.

Lei, Y., Huang, Y., Wen, X., Yin, Z., Zhang, Z., and Klionsky, D. J. (2022). How Cells Deal with the Fluctuating Environment: Autophagy Regulation under Stress in Yeast and Mammalian Systems. Antioxidants (Basel) 11. doi:10.3390/antiox11020304.

Levine, B., and Kroemer, G. (2008). Autophagy in the pathogenesis of disease. Cell 132, 27–42. doi:10.1016/j.cell.2007.12.018.

Li, G., Poulsen, M., Fenyvuesvolgyi, C., Yashiroda, Y., Yoshida, M., Simard, J. M., Gallo, R. C., and Zhao, R. Y. (2017). Characterization of cytopathic factors through genome-wide analysis of the Zika viral proteins in fission yeast. Proc. Natl. Acad. Sci. USA 114, E376–E385. doi:10.1073/pnas.1619735114.

Liu, D. X., Fung, T. S., Chong, K. K.-L., Shukla, A., and Hilgenfeld, R. (2014). Accessory proteins of SARS-CoV and other coronaviruses. Antiviral Res. 109, 97–109. doi:10.1016/j.antiviral.2014.06.013.

Macreadie, I. G., Castelli, L. A., Hewish, D. R., Kirkpatrick, A., Ward, A. C., and Azad, A. A. (1995). A domain of human immunodeficiency virus type 1 Vpr containing repeated H(S/F)RIG amino acid motifs causes cell growth arrest and structural defects. Proc. Natl. Acad. Sci. USA 92, 2770–2774. doi:10.1073/pnas.92.7.2770.

Mao, J., Lin, E., He, L., Yu, J., Tan, P., and Zhou, Y. (2019). Autophagy and viral infection. Adv. Exp. Med. Biol. 1209, 55–78. doi:10.1007/978-981-15-0606-2_5.

McBride, R., and Fielding, B. C. (2012). The role of severe acute respiratory syndrome (SARS)-coronavirus accessory proteins in virus pathogenesis. Viruses 4, 2902–2923. doi:10.3390/v4112902.

Miao, G., Zhao, H., Li, Y., Ji, M., Chen, Y., Shi, Y., Bi, Y., Wang, P., and Zhang, H. (2021). ORF3a of the COVID-19 virus SARS-CoV-2 blocks HOPS complex-mediated assembly of the SNARE complex required for autolysosome formation. Dev. Cell 56, 427–442.e5. doi:10.1016/j.devcel.2020.12.010.

Miller, K., McGrath, M. E., Hu, Z., Ariannejad, S., Weston, S., Frieman, M., and Jackson, W. T. (2020). Coronavirus interactions with the cellular autophagy machinery. Autophagy 16, 2131–2139. doi:10.1080/15548627.2020.1817280.

Nakatogawa, H. (2020). Mechanisms governing autophagosome biogenesis. Nat. Rev. Mol. Cell Biol. 21, 439–458. doi:10.1038/s41580-020-0241-0.

Nieva, J. L., Madan, V., and Carrasco, L. (2012). Viroporins: structure and biological functions. Nat. Rev. Microbiol. 10, 563–574. doi:10.1038/nrmicro2820.

Park, H.-S., Lee, S. C., Cardenas, M. E., and Heitman, J. (2019). Calcium-Calmodulin-Calcineurin Signaling: A Globally Conserved Virulence Cascade in Eukaryotic Microbial Pathogens. Cell Host Microbe 26, 453–462. doi:10.1016/j.chom.2019.08.004.

Pradel, B., Robert-Hebmann, V., and Espert, L. (2020). Regulation of innate immune responses by autophagy: A goldmine for viruses. Front. Immunol. 11, 578038. doi:10.3389/fimmu.2020.578038.

Qu, Y., Wang, X., Zhu, Y., Wang, W., Wang, Y., Hu, G., Liu, C., Li, J., Ren, S., Xiao, M. Z. X., et al. (2021). ORF3a-Mediated Incomplete Autophagy Facilitates Severe Acute Respiratory Syndrome Coronavirus-2 Replication. Front. Cell Dev. Biol. 9, 716208. doi:10.3389/fcell.2021.716208.

Reggiori, F., Monastyrska, I., Verheije, M. H., Calì, T., Ulasli, M., Bianchi, S., Bernasconi, R., de Haan, C. A. M., and Molinari, M. (2010). Coronaviruses Hijack the LC3-I-positive EDEMosomes, ER-derived vesicles exporting short-lived ERAD regulators, for replication. Cell Host Microbe 7, 500–508. doi:10.1016/j.chom.2010.05.013.

Ren, Y., Shu, T., Wu, D., Mu, J., Wang, C., Huang, M., Han, Y., Zhang, X.-Y., Zhou, W., Qiu, Y., et al. (2020). The ORF3a protein of SARS-CoV-2 induces apoptosis in cells. Cell Mol Immunol 17, 881–883. doi:10.1038/s41423-020-0485-9.

Richetta, C., and Faure, M. (2013). Autophagy in antiviral innate immunity. Cell Microbiol. 15, 368–376. doi:10.1111/cmi.12043.

Rienzo, A., Pascual-Ahuir, A., and Proft, M. (2012). The use of a real-time luciferase assay to quantify gene expression dynamics in the living yeast cell. Yeast 29, 219–231. doi:10.1002/yea.2905.

Sanders, J. M., Monogue, M. L., Jodlowski, T. Z., and Cutrell, J. B. (2020). Pharmacologic Treatments for Coronavirus Disease 2019 (COVID-19): A Review. JAMA 323, 1824–1836. doi:10.1001/jama.2020.6019.

Shroff, A., and Nazarko, T. Y. (2021). The Molecular Interplay between Human Coronaviruses and Autophagy. Cells 10. doi:10.3390/cells10082022.

Solvik, T. A., Nguyen, T. A., Tony Lin, Y.-H., Marsh, T., Huang, E. J., Wiita, A. P., Debnath, J., and Leidal, A. M. (2022). Secretory autophagy maintains proteostasis upon lysosome inhibition. J. Cell Biol. 221. doi:10.1083/jcb.202110151.

Stathopoulos, A. M., and Cyert, M. S. (1997). Calcineurin acts through the CRZ1/TCN1-encoded transcription factor to regulate gene expression in yeast. Genes Dev. 11, 3432–3444. doi:10.1101/gad.11.24.3432.

Swanton, C., and Jones, N. (2001). Strategies in subversion: de-regulation of the mammalian cell cycle by viral gene products. Int J Exp Pathol 82, 3–13. doi:10.1046/j.1365-2613.2001.00165.x.

Thewes, S. (2014). Calcineurin-Crz1 signaling in lower eukaryotes. Eukaryotic Cell 13, 694–705. doi:10.1128/EC.00038-14.

Tokumitsu, H., and Sakagami, H. (2022). Molecular Mechanisms Underlying Ca2+/Calmodulin-Dependent Protein Kinase Kinase Signal Transduction. Int. J. Mol. Sci. 23. doi:10.3390/ijms231911025.

Wolff, G., Melia, C. E., Snijder, E. J., and Bárcena, M. (2020). Double-Membrane Vesicles as Platforms for Viral Replication. Trends Microbiol. 28, 1022–1033. doi:10.1016/j.tim.2020.05.009.

Wu, F., Zhao, S., Yu, B., Chen, Y.-M., Wang, W., Song, Z.-G., Hu, Y., Tao, Z.-W., Tian, J.-H., Pei, Y.-Y., et al. (2020). A new coronavirus associated with human respiratory disease in China. Nature 579, 265–269. doi:10.1038/s41586-020-2008-3.

Young, C. L., Raden, D. L., and Robinson, A. S. (2013). Analysis of ER resident proteins in Saccharomyces cerevisiae: implementation of H/KDEL retrieval sequences. Traffic 14, 365–381. doi:10.1111/tra.12041.

Zhang, J., Cruz-Cosme, R., Zhuang, M.-W., Liu, D., Liu, Y., Teng, S., Wang, P.-H., and Tang, Q. (2020). A systemic and molecular study of subcellular localization of SARS-CoV-2 proteins. Signal Transduct. Target. Ther. 5, 269. doi:10.1038/s41392-020-00372-8.

Zhang, Y., Sun, H., Pei, R., Mao, B., Zhao, Z., Li, H., Lin, Y., and Lu, K. (2021). The SARS-CoV-2 protein ORF3a inhibits fusion of autophagosomes with lysosomes. Cell Discov. 7, 31. doi:10.1038/s41421-021-00268-z.

Zhang, Y.-Z., and Holmes, E. C. (2020). A Genomic Perspective on the Origin and Emergence of SARS-CoV-2. Cell 181, 223–227. doi:10.1016/j.cell.2020.03.035.

Zhao, R. Y. (2017). Yeast for virus research. Microb. Cell 4, 311–330. doi:10.15698/mic2017.10.592.

Zhao, Y., Cao, J., O’Gorman, M. R., Yu, M., and Yogev, R. (1996). Effect of human immunodeficiency virus type 1 protein R (vpr) gene expression on basic cellular function of fission yeast Schizosaccharomyces pombe. J. Virol. 70, 5821–5826. doi:10.1128/JVI.70.9.5821-5826.1996.

Zhao, Z., Lu, K., Mao, B., Liu, S., Trilling, M., Huang, A., Lu, M., and Lin, Y. (2021). The interplay between emerging human coronavirus infections and autophagy. Emerg. Microbes Infect. 10, 196–205. doi:10.1080/22221751.2021.1872353.

Zmasek, C. M., Lefkowitz, E. J., Niewiadomska, A., and Scheuermann, R. H. (2022). Genomic evolution of the Coronaviridae family. Virology 570, 123–133. doi:10.1016/j.virol.2022.03.005.

